# Detecting *CYP2C19* deletions from genotyping array signals using neural networks

**DOI:** 10.64898/2026.08.21.746170

**Authors:** Burak Yelmen, Robin J. Hofmeister, Viido Kaur Lutsar, Marc Finianos, Benjamin C. Stone, Maarja Jõeloo, Kristi Krebs, Paula Ann Kivistik, Steven Smit, Estonian Biobank Research Team, Mait Metspalu, Georgi Hudjashov, Lili Milani

## Abstract

Since copy number variations (CNVs) in pharmacogenes can cause significant alterations in drug metabolism, their reliable detection is of high importance both for large-scale studies and personalized medicine. Whole-genome sequencing, and specifically long-read sequencing, is the gold standard for CNV detection. Despite increasing availability of these technologies, genotyping arrays are still widely used as cost-effective alternatives in biobank and clinical settings, yet calling CNVs based on array intensity signals is challenging due to low base pair resolution. In this work, we developed a neural network model, nnCNV, to predict deletions in the *CYP2C19* pharmacogene region from array intensity signals. We compared our method to the most widely used algorithm, PennCNV, and demonstrated better performance reaching 100% accuracy in the test dataset. Furthermore, we predicted probe-by-probe *CYP2C19* deletion coordinates for all Estonian Biobank samples using nnCNV and PennCNV, and validated these predictions using an identity-by-descent (IBD) sharing method, which also demonstrated superior nnCNV performance. For the deletion samples with conflicting PennCNV and nnCNV predictions, we performed PCR analysis for validation, which showed 97% precision for nnCNV compared to 23% for PennCNV. Finally, we assessed the gradient-based feature importance maps and showed that nnCNV utilizes signal intensity information not only from deletion probes, but also from probes in flanking regions. Our results demonstrate that long-range information, which cannot be utilized by hidden Markov models, can improve CNV calling.

## Introduction

Copy number variations (CNVs) are a major class of structural variation (SV) in the human genome with high biological and clinical relevance: they can disrupt coding sequences and perturb regulatory elements, thereby altering various phenotypic outcomes [1]. CNVs are especially of interest in the context of pharmacogenes due to their ability to alter drug metabolism drastically, hence, their reliable detection is of high importance both for large-scale pharmacogenomics studies and personalized medical interventions [2, 3]. One such important pharmacogene is *CYP2C19*, which codes for the CYP2C19 enzyme. CYP2C19 is part of the CYP450 superfamily involved in the metabolism of multiple clinically-relevant drugs such as clopidogrel, voriconazole, proton pump inhibitors and antidepressants [4]. Sequencing the genomes of hundreds of individuals from different populations has highlighted the significance of SVs in *CYP2C19* [4]. The Pharmacogene Variation Consortium (PharmVar) [5] now lists star alleles for complete gene deletions (*CYP2C19*36)*, partial deletions involving upstream regions and at least exon 1 (*CYP2C19*37*), and partial deletions involving exons 2-5 (*CYP2C19*42)* (S1 Fig) with cumulative frequencies up to 2% reported in the Estonian and Finnish populations [6, 7].

Most reliable and high resolution detection of SVs can be achieved with whole-genome sequencing (WGS) and more recently long-read sequencing technologies [8, 9]. Despite the decreased costs and increased availability of WGS, genotyping arrays are still widely used as cost-efficient alternatives to capture genome-wide variation. These arrays are designed based on a specific set of probes to represent highly variable or functionally relevant single nucleotide variants (SNVs) [10]. For each marker, fluorescence probe intensities are normalized for the A and B alleles and summarized as total signal intensity and allelic balance. Log R ratio (LRR) is the log2 ratio of observed to expected normalized total intensity and provides a measure of relative DNA copy number. B-allele frequency (BAF) is an intensity-derived allelic ratio, calibrated from the relative A/B signal using canonical genotype clusters, with expected values near 0, 0.5 and 1 for AA, AB, and BB genotypes, respectively. Together, LRR and BAF provide complementary information for CNV detection, with LRR capturing total copy number and BAF capturing the relative proportion of the two alleles.

One of the most widely used algorithms for CNV detection from array signals is PennCNV, which is based on a hidden Markov model (HMM) to infer copy number states from the observed signal intensity values (i.e., BAF and LRR) [11]. Despite its widespread use, multiple comparative studies, while highlighting PennCNV as one of the best algorithms, reported modest precision and accuracy performance [12, 13]. One potential explanation for this could be an inherent limitation of HMM-based algorithms: copy number state at each SNP is assumed to depend only on the copy number state of the previous SNP, limiting their capacity to exploit long-range information.

In this work, we developed a neural network model, nnCNV, to predict deletion status for each probe in the *CYP2C19* region using BAF and LRR signals. We first utilized a subset of the Estonian Biobank (EstBB) [14] individuals with both genotyping and long-read sequencing data to train the model and to assess performance in comparison to PennCNV. We then predicted the deletion status for all biobank samples, and validated these predictions using an approach based on identity-by-descent (IBD) sharing between relatives. Our findings demonstrated that nnCNV either matches or outperforms PennCNV in all performance metrics. To investigate the roots of this performance gain, we analyzed the gradient-based feature attribution maps of nnCNV and demonstrated that multiple probes over the whole target region had high scores, suggesting that long-range signal information from flanking regions can be useful for predictions.

## Materials and Methods

### Dataset

We obtained LRR and BAF signals from four different GSA array versions (Illumina’s GSA-MD-24v1, GSA-MD-24v2, ESTchip1 GSA-MD-24v2, ESTchip2 GSA-MD-24v3) over 211,257 individuals from the EstBB [14]. For 5095 individuals, corresponding PacBio long-read sequencing data was also available, which was used to obtain ground-truth CNV labels using Sawfish [15] for the *CYP2C19* region. Five samples with potential contamination during sequencing and predicted to have *CYP2C19* deletions by Sawfish were removed in subsequent analyses.

### SV calling with Sawfish and PennCNV

SV calling was performed using 20x long-read genomes generated with PacBio HiFi technology. Raw sequencing data was mapped using pbmm2 v1.17.0 and HiFi preset. The SVs were detected with Sawfish v2.2.1 using default parameters.

PennCNV v1.0.4 [11] was run with default parameters and GC correction in 17 genotyping batches. Prior to CNV detection, we removed (1) custom EstChip content added to GSA arrays, (2) multiallelic probes, (3) CNV probes, and (4) duplicated probes. For each batch, we generated a batch-specific population B allele frequency file. After the exclusion of duplicates and 1,728 samples with inflated intensity profiles, we retained raw calls for a total of 210,213 unique samples. We defined carriers of the *CYP2C19* deletion as samples with previously established CNV breakpoints within chr10: 96,497,325-96,559,109 (hg19). Carriers of other deletion or duplication calls overlapping CYP2C19 were flagged as ambiguous and excluded from further analyses.

The final feature set (i.e., genotyping array probe positions with BAF and LRR values) to be used by the neural network model was determined based on Sawfish and PennCNV predictions. More specifically, the lowest and the highest base pair positions were combined from Sawfish and PennCNV predictions over 5090 individuals, and ±500 kb was added to the terminal upstream and downstream positions to obtain the final border points, which resulted in 410 probe positions spanning over the *CYP2C19* region.

### Confirmation of Sawfish predictions

Sawfish detected deletions in 165 samples, with the most common being the CYP2C19*37 variant (21-probe-length deletion) in 141 samples. The other variants were exon2-5 long (13-probe-length deletion, 3 samples), exon2-5 short (12-probe-length deletion, 9 samples) - both now designated CYP2C19*42, and small del (0-probe-length deletion as there was no probe in the deletion region, 12 samples). To further confirm these deletions and mitigate potential false positives, we assessed the aligned reads for samples with deletions. Specifically, the BAM alignment file was first subset with samtools v1.22.1 to retain alignments spanning *CYP2C19* and the putative deletion region [16]. Deletions and insertions ≥50 bp were extracted from the CIGAR-string using Pysam v0.23.3 [17, 18]. To detect larger structural deletions, alignments were screened for same-orientation supplementary alignments. Their break end (BND) coordinates were recorded, deletion size was inferred from the genomic gap, and the read depth was quantified on the 1-bp inner borders. The total amount of reads supporting the BND was compared to the corresponding read count in that region. For all Sawfish deletion carriers, at least 13% of the reads supported the presence of a deletion, with mean support being 47%. This is consistent with mostly heterozygous deletions carriers, as our dataset had only two homozygous individuals.

### Neural network model

The prediction model, which we name nnCNV, is a feedforward fully connected neural network for predicting the deletion status of each probe position for the 410-probe-length *CYP2C19* region. For model development, we first divided the dataset of 5090 samples into train/validation and test sets with a 0.80-0.20 split. Since some deletion patterns were rare, the split was done so that each deletion pattern would be also present in the test set. The train/validation set was further split into train and validation sets (0.90-0.10 split), which were used for model optimization and training. The test set was not used until the final model assessment.

The architecture and hyperparameter search was done using 4-fold cross-validation over 36 configurations (hidden layer size = 205, 410, 820, 1640; learning rate = 1e-4, 5e-5, 1e-5; dropout = 0.1, 0.25, 0.5) each repeated 2 times using train/validation set. The configuration with the lowest mean validation accuracy was selected as the best configuration. The final architecture consists of a linear layer with input size 820 (for concatenated BAF and LRR values) and output size 1640; a dropout layer (p = 0.5) followed by ReLU activation; a linear layer with input size 1640 and output size 1640; a dropout layer (p = 0.5) followed by ReLU activation; a linear layer with input size 1640 and output size 820 (corresponding to two logits for each class prediction). Based on this architecture, the total trainable parameter size was 5,383,300.

The training was performed using cross-entropy loss, Adam optimizer with learning rate = 5e-5, and batch size = 128. The sampling was done so that each batch had an equal number of samples with (n = 64) and without (n = 64) deletion. Training was stopped using early stopping with 200 epoch patience (i.e., training stopped if the validation loss did not decrease over 200 epochs), and the best model over all epochs was saved as the final model. nnCNN was trained up to 1328 epochs and best model weights based on the validation loss was obtained from epoch 1128 (S2 Fig). The model was coded with Python-3.9 and PyThorch-1.11 [19].

### Biobank-scale detection and validation

After model training and performance evaluation, we predicted the deletion status of the *CYP2C19* region of all 211,257 EstBB individuals using nnCNV and PennCNV. To assess the validity of these predictions, we estimated CNV inheritance rates (R) by combining identity-by-descent (IBD) and CNV-sharing information between carriers and their close relatives. IBD was inferred using THORIN [20], and individuals were classified as IBD0, IBD1, or IBD2 based on haplotype-sharing probabilities averaged within a 1 Mb window around each CNV, thereby capturing whether the SNV haplotypic background of the CNV is shared identical by descent. We considered a haplotype to be shared IBD when the average per-variant IBD probability in this window was ≥ 0.75. In parallel, CNVs observed in relatives were classified as shared or not shared based on reciprocal overlap, with CNVs considered shared when the overlap was ≥ 0.75. We nevertheless also assessed different IBD and CNV sharing cutoffs (0.1, 0.25, 0.5, 0.75, 0.95) in comparisons. By combining IBD status and CNV sharing, we identified configurations consistent with inherited versus de novo events. We then derived a conservative estimate of the CNV inheritance rate across all informative relative pairs, prioritizing the most informative relationships when multiple relatives were available, and estimated standard errors using parametric bootstrap based on binomial sampling.

### PCR validation

To further assess the validation of predictions from the whole biobank dataset, we compiled a subset of samples which are **(i)** predicted to have a deletion by PennCNV and no deletion by nnCNV (n = 173), **(ii)** predicted to have a deletion by nnCNV and no deletion by PennCNV (n = 207), and **(iii)** predicted to have unique patterns by nnCNV not matching Sawfish patterns (n = 8). These samples with deletions were then validated with PCR. 35 cycles of PCR amplifications from genomic DNA were carried out using HOT FIREPol GC Master Mix reagents (Solis BioDyne OÜ) and deletion specific primers with annealing temperature of 61 ^*◦*^C [7]. Primers and their genomic coordinates are listed in the S1 Table. PCR products were visualised on 1.5% TBE agarose gel. After removing the samples for which PCR did not work reliably (n = 18), the total sample size was 370.

### Data availability

Code for the model training and evaluation is available at https://github.com/genodeco/CYP2C19-CNV-detection with mock data. Access to the Estonian Biobank dataset is subject to an application procedure, for which details can be found at https://genomics.ut.ee/en/content/estonian-biobank.

## Results

### Model assessment

Following model optimization and training, we first assessed nnCNV predictions on the test data and compared them to PennCNV based on position-level predictions (i.e., probe-by-probe predictions, Fig 1a) and sample-level predictions (i.e., binary deletion or no deletion for each sample, Fig 1b). Overall, both models performed well but nnCNV performance was superior with 100% accuracy. The performance difference was especially apparent for position-level assessment since PennCNV was prone to false negatives: it failed to predict 106 out of 625 total deletion positions and 1 out of 31 total individuals with deletions.

**Figure 1.**
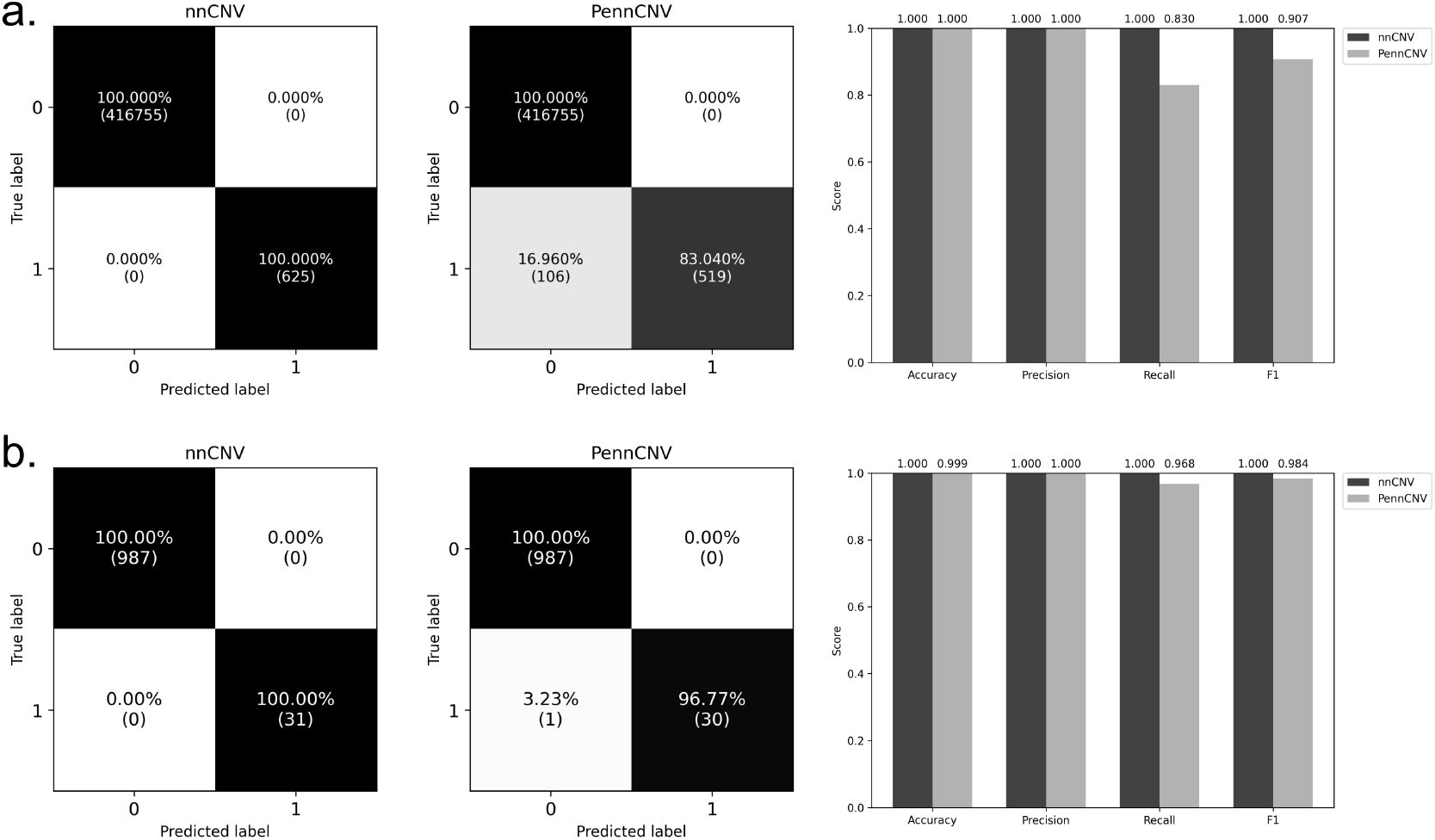
Confusion matrices and performance metrics of nnCNV and PennCNV over test dataset based on **a)** position-level predictions (i.e., probe-by-probe prediction performance based on long-read sequencing labels) and **b)** sample-level predictions (i.e., binary deletion or no deletion prediction for each sample).

### Biobank-scale prediction

After performance analysis on the test data, we predicted *CYP2C19* region CNVs for all EstBB samples using nnCNV and PennCNV, and compared the results. PennCNV detected deletions in 4013 individuals and nnCNV in 4047 individuals. We compared the combined set of individuals detected to have deletions (4220) between these two models in terms of variability and agreement (Fig 2). PennCNV showed high variance in the deletion region over individuals whereas for nnCNV, the deletion region borders were well defined, matching the break points of the deletions in the training samples (Fig 2a). Sample-level agreement was generally high between the two models but 207 samples were predicted to have no deletions by PennCNV while nnCNV predicted deletions, and 173 samples were predicted to have deletions by PennCNV while nnCNV predicted no deletions. There was higher discrepancy in terms of position-level agreement, especially for positive predictions since ∼ 25% of positions predicted to be deletions by nnCNV were predicted to be no deletion by PennCNV (Fig 2b).

**Figure 2.**
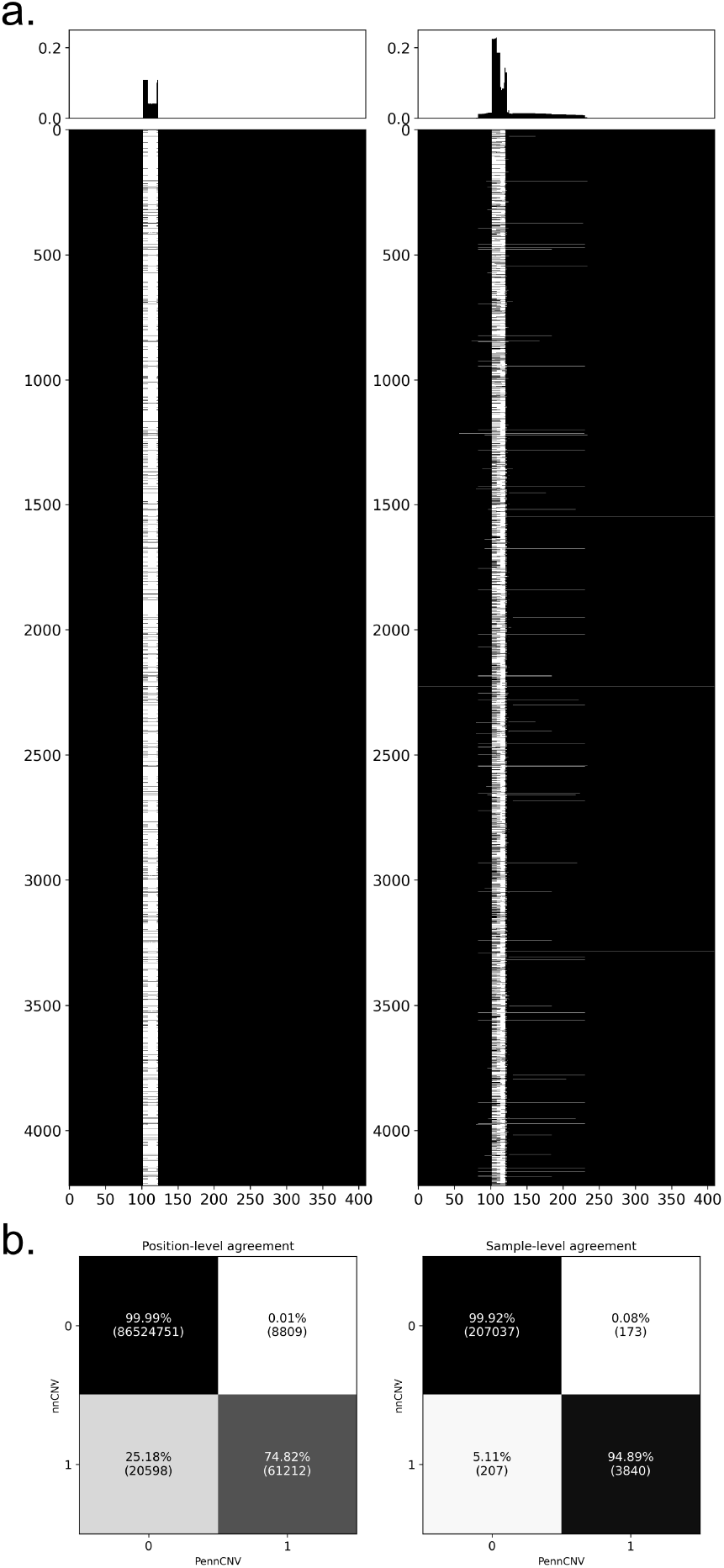
Biobank-scale prediction of *CYP2C19* deletions. **a)** Comparison of nnCNV (left) and PennCNV right) for individuals (y-axis) with deletions over target region probes (x-axis). White color indicates deletion and black color no deletion. Plots above demonstrate variance over probe positions. Position-level (left) and sample-level (right) agreement matrices of PennCNV (x-axis) and nnCNV (y-axis).

Since our test data was limited in terms of samples with deletions and we did not have whole-genome sequencing data for most of the EstBB samples, to assess the validity of the predictions over all the biobank samples, we analyzed CNV inheritance rates (R, see Materials and Methods). Essentially, R allows us to infer the reliability of CNV predictions by validating whether they follow expectations based on IBD sharing among close relatives. For all different cutoff thresholds, nnCNV achieved R between ∼ 0.9 and ∼ 1.0 whereas PennCNV R ranged from ∼ 0.5 to ∼ 0.7, indicating that nnCNV predictions were more consistent with expected inheritance patterns compared to PennCNV (Fig 3). In addition, we compared the performances for different quality score (QS) thresholds for PennCNV (S3 Fig). Filtering with a QS threshold of 0.1 slightly improved PennCNV performance, but stricter thresholds reduced the sample sizes substantially (2517, 1276, 83, 6 samples for QS = 0, 0.1, 0.25, 0.5, respectively) and increased standard deviation.

**Figure 3.**
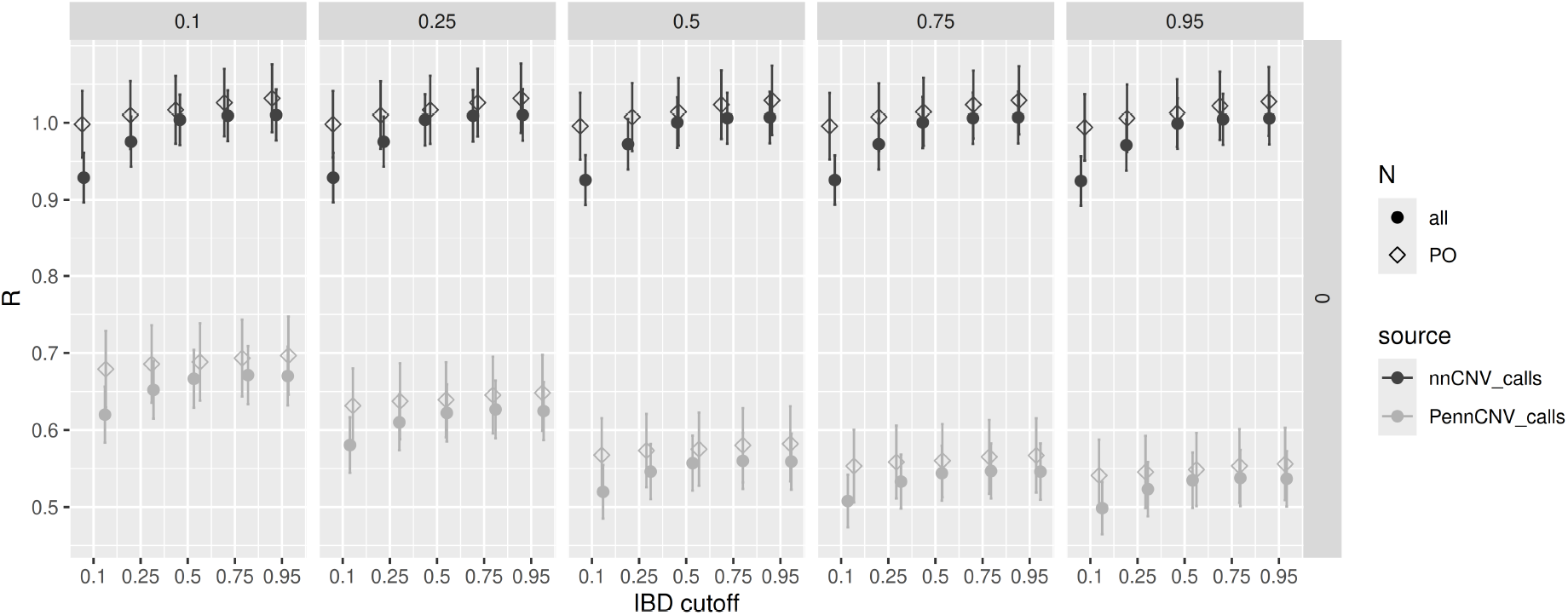
Comparison of CNV inheritance rate R (y-axis) between nnCNV and PennCNV over different IBD sharing (x-axis) and CNV sharing (different columns) cutoffs. Calculations based on all samples (all) and only on parent-offspring trios (PO) are indicated with different shapes. Vertical lines indicate standard errors obtained with bootstrap.

### PCR validation

We further assessed the conflicting PennCNV and nnCNV deletion predictions over all biobank samples using PCR (see Materials and Methods). Among the samples predicted to have no deletion by PennCNV and deletion by nnCNV (n=195), 175 showed a deletion signal by PCR analysis (false negative PennCNV signals). Out of the remaining 20 samples, 5 samples showed no deletion signals (false positive nnCNV signals), and at least one deletion-specific primer did not work reliably for the other 15 samples with inconclusive results.

Among the samples predicted to have a deletion by PennCNV and no deletion by nnCNV (n=167), 30 showed a deletion signal by PCR analysis (false negative nnCNV signals). Out of the remaining 137 samples, 98 samples showed no deletion signals (false positive PennCNV signals), and at least one deletion-specific primer did not work reliably for the other 48 samples.

Overall, for this collection of samples with conflicting predictions, precision was 97% for nnCNV and 23% for PennCNV. Confusion matrices summarizing these findings can be seen in S4 Fig.

For the 8 samples predicted to have a unique deletion pattern by nnCNV, 7 showed deletion signal in PCR based on 3 primers, and 1 showed no deletion signal. All of these 8 samples were also predicted to have deletions by PennCNV.

### Gradient-based feature attribution maps

To investigate the performance gain of nnCNV over PennCNV, we computed the Jacobian of the model outputs with respect to BAF and LRR input features for each sample, and visualized the resulting input-output sensitivity matrices (Fig 4, S5 Fig). Interestingly, gradient attribution maps demonstrated that even though features at the specific deletion region tended to have the highest absolute attribution scores, BAF and LRR input features over the whole 410 probe region were utilized for predictions. In addition, LRR features had higher scores than BAF features, suggesting that LRR is more relevant for detecting deletions as expected [11, 21].

**Figure 4.**
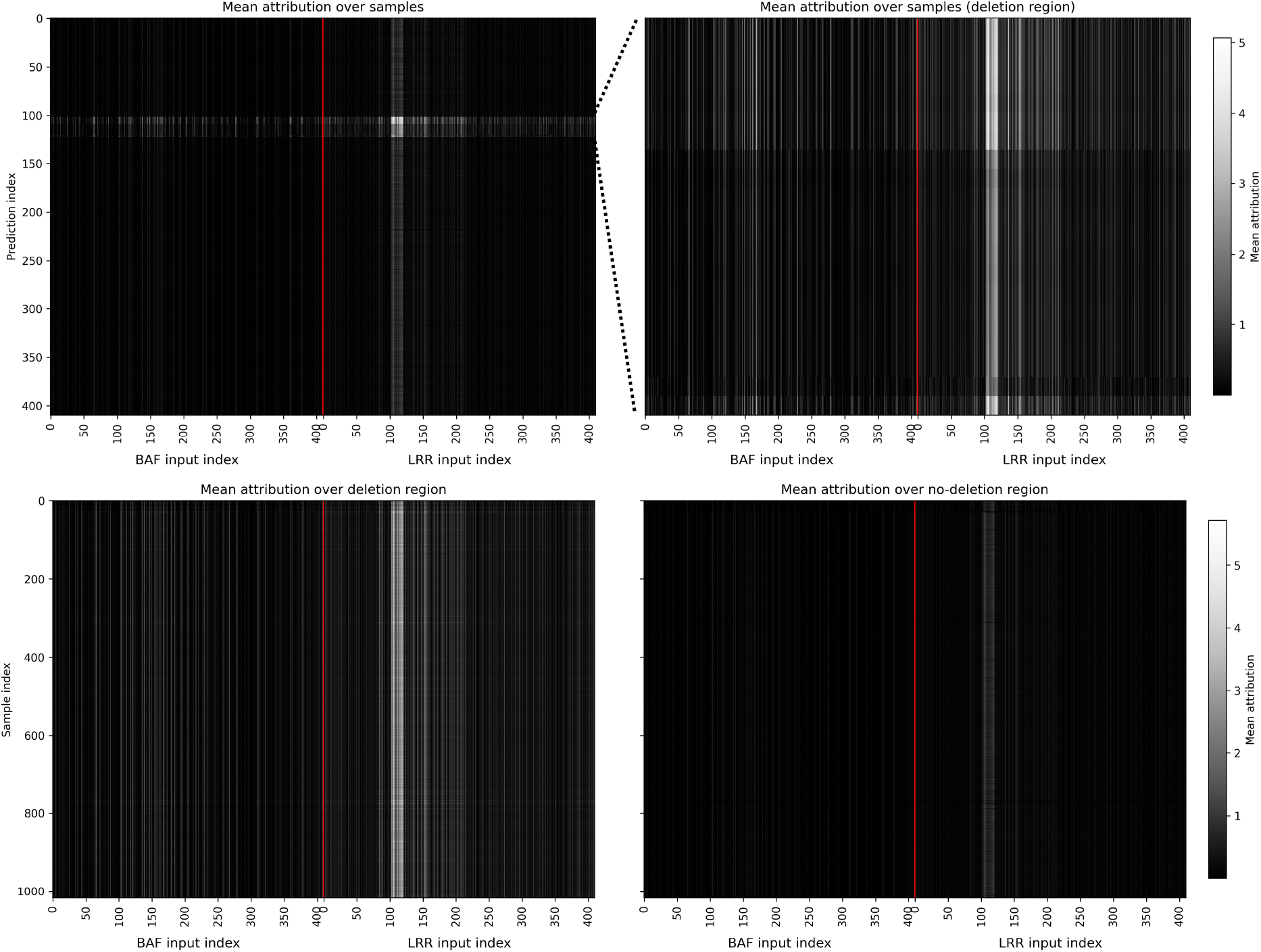
Gradient-based feature attribution heatmaps for nnCNV using test samples. **a)** Sample-averaged attribution matrix showing the gradient-based attribution of each input feature position (x-axis) to each output node/position (y-axis). **b)** Target-region-averaged attribution heatmaps for the deletion region (left) and no-deletion region (right), showing attribution scores across input feature positions (x-axis) for individual samples (y-axis). True deletion region spans probes 102-122. Vertical red lines separate BAF and LRR input indices.

## Discussion

In this work, we developed a neural network model, nnCNV, to detect deletions in the *CYP2C19* region. Our results based on the test data and over the whole biobank samples demonstrated superior performance for nnCNV compared to the HMM-based PennCNV algorithm. We further assessed the conflicting calls by PennCNV and nnCNV via PCR analysis, and demonstrated again substantially better performance for nnCNV (S4 Fig). A potential explanation for this is the ability of the fully-connected neural networks to capture long-range information. Indeed, we were able to confirm this by assessing the gradient-based feature importance maps (Fig 4, S3 Fig). We further investigated the correlation patterns for LRR and BAF over the region, and found that there was correlation between probe positions even outside the deletion borders (S6 Fig). However, it is not trivial to assess whether there are additional CNV-related signals outside deletion borders (i.e., signals which carry unique predictive information not available in the deletion probes) or whether correlations simply indicate duplicated information due to the array framework, normalization or haplotype structure. Despite this nuance, our results demonstrate that long-range information, which cannot be utilized by HMM-based models, can be useful in CNV calling. An important limitation of our approach is the locus specificity. Unlike PennCNV, nnCNV has only been optimized/validated for the *CYP2C19* region in its current configuration, and new models need to be trained and potentially optimized for other regions with different deletion and duplication compositions. We chose this region due to its pharmacological importance [6] and as a proof-of-concept application due to relatively easy-to-characterize CNV variation consisting only of heterozygous (except for 2 samples which were homozygous) deletions. This simplicity allowed us to have an architecture for binary prediction for each probe position, without considering duplication or heterozygous/homozygous status. It remains to be seen whether superior neural network performance can be achieved over the whole genome or other specific loci with more difficult CNV patterns.

An interesting finding was the 8 different deletion patterns detected by nnCNV, which were not present in the training and test data. We confirmed that 7 of these had some type of deletion via PCR, which suggests that most of these patterns are likely partially mis-called versions of the already characterized deletion variants. However, since nnCNV is trained for probe-by-probe prediction instead of categorical prediction for the whole region, it could potentially detect novel rare deletion variants. Since these novel variants might require specific primers for PCR detection, the most reliable way to map their exact coordinates would be long-read sequencing.

Despite the increasing availability of WGS owing to reduced costs and new technologies, genotyping arrays are likely to persist in both research and clinical settings as cheaper alternatives suitable for fast, large-scale analyses. In this context, recent studies have demonstrated that machine learning and deep learning approaches may improve the accuracy of CNV detection and validation [22–24]. Our findings also demonstrate that neural networks can be used to call gene-specific CNVs with high precision and fewer false negatives than PennCNV. After experimental and external validation, nnCNV can be considered a reliable alternative or a complementary method to PennCNV for CNV calling in the *CYP2C19* region when WGS is not available.

## Author Contributions

Conceptualization: BY, LM. Data curation: BY, KK, EBRT, GH. Methodology: BY, RJH, MF. Software: BY, VKL. Investigation: BY, RJH, VKL, MF, BCS, MJ, KK, PAK, SS, GH. Formal analysis: BY, RJH, VKL. Visualization: BY, RJH. Supervision: BY, MM, LM. Project administration: BY. Funding acquisition: BY, MM, LM. Original draft: BY. Review & editing: BY, RJH, MF, MJ, PAK, GH, LM.

## Acknowledgements

Thanks to the High Performance Computing Center (HPC) of the University of Tartu for providing computational resources. Thanks to the Estonian Biobank Research Team (Andres Metspalu, Lili Milani, Tõnu Esko, Reedik Mägi, Mari Nelis, Georgi Hudjashov) for the genotype and phenotype datasets used in this work. BY was supported by the Estonian Research Council grant PSG1179. LM was supported by the Estonian Research Council grant PRG2625. BCS was supported by the U.S. Fulbright Program. The research was conducted using the Estonian Center of Genomics/Roadmap II funded by the Estonian Research Council (project number TT17). The research was conducted using the research infrastructure “Estonian Center for Genomics” funded by the Estonian Research Council (project number TARISTU24-TK19). This work has received funding from the European Union’s Horizon Europe research and innovation programme under grant agreement No 101060011. Views and opinions expressed are however those of the author(s) only and do not necessarily reflect those of the European Union or European Research Executive Agency.

## Ethics statement

The activities of the EstBB are regulated by the Human Genes Research Act, which was adopted in 2000 specifically for the operations of the EstBB. Individual level data analysis was carried out under ethical approval 1.1-12/624 (issued 24.03.2020), 1.1-12/2618 (issued 04.08.2022), 1.1-12/3454 (issued 20.10.2022) and 1.1-12/906 (issued 31.03.2026) from the Estonian Committee on Bioethics and Human Research (Estonian Ministry of Social Affairs).

