## Supplementary Figures for "Detecting *CYP2C19* deletions from genotyping array signals using neural networks"

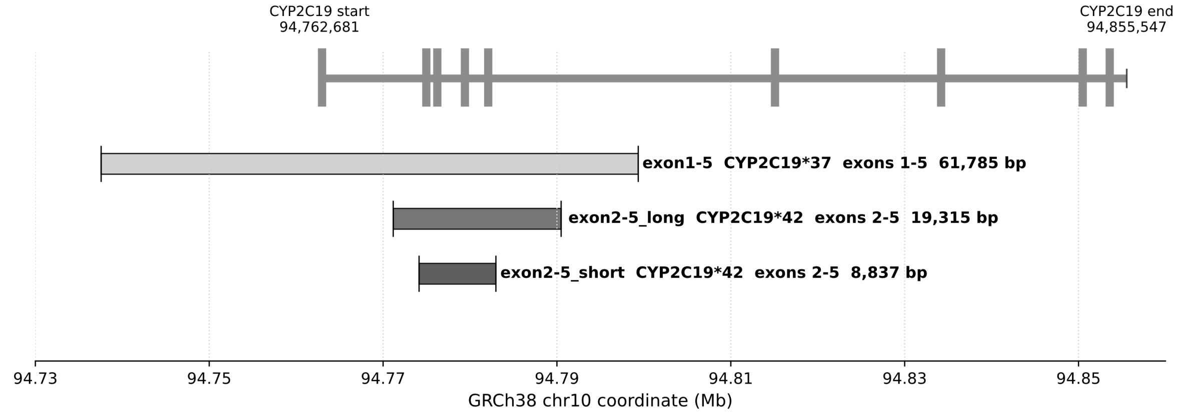


**S1 Fig.** Schematic of *CYP2C19* deletion coordinates.

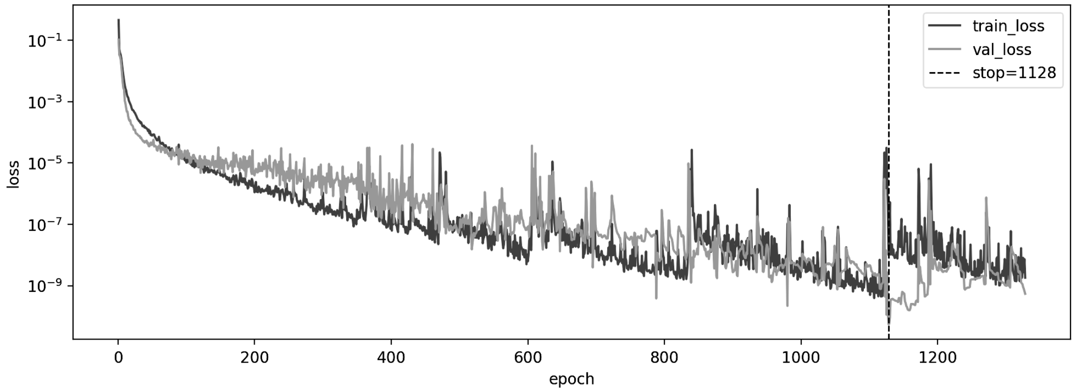


**S2 Fig.** Loss plot of nnCNV training. Vertical dashed line shows the lowest validation loss epoch, 1128.

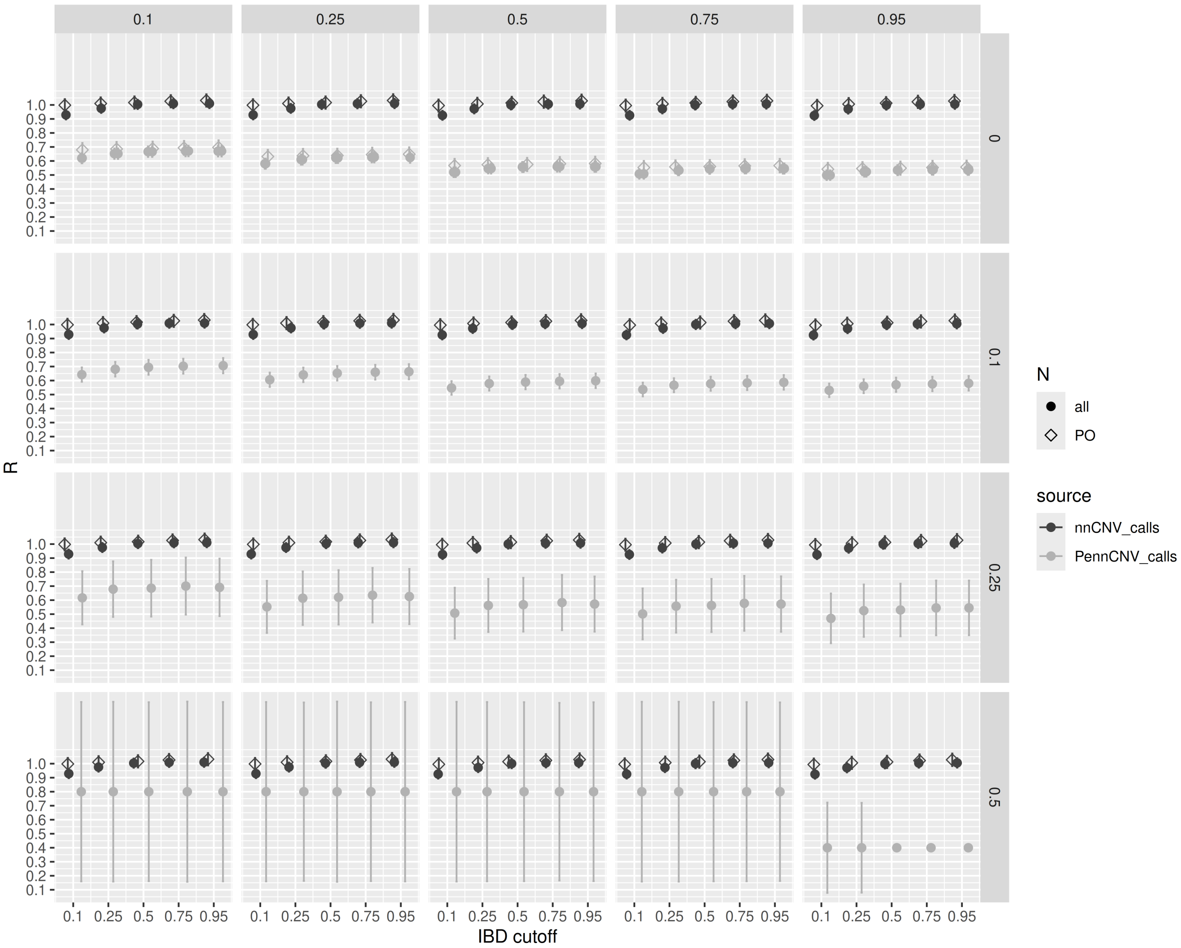


**S3 Fig.** Comparison of CNV inheritance rate R (y-axis) between nnCNV and PennCNV over different IBD sharing (x-axis) and CNV sharing (different columns) cutoffs. Each row is a different quality score cutoff for PennCNV. Calculations based on all samples (all) and only on parent-offspring trios (PO) are indicated with different shapes. Vertical lines indicate standard errors obtained with bootstrap.

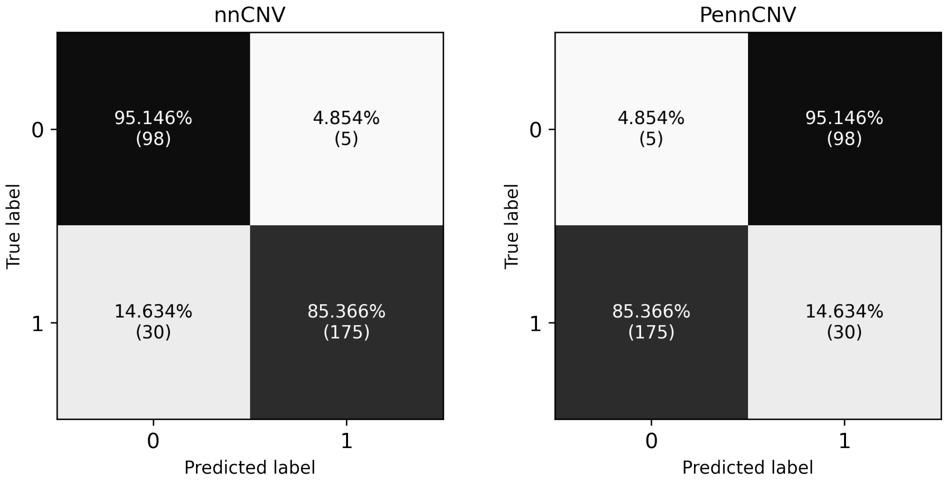


**S4 Fig.** Confusion matrices for nnCNV and PennCNV based on PCR validation of samples predicted to have deletion in one method and no deletion in the other.

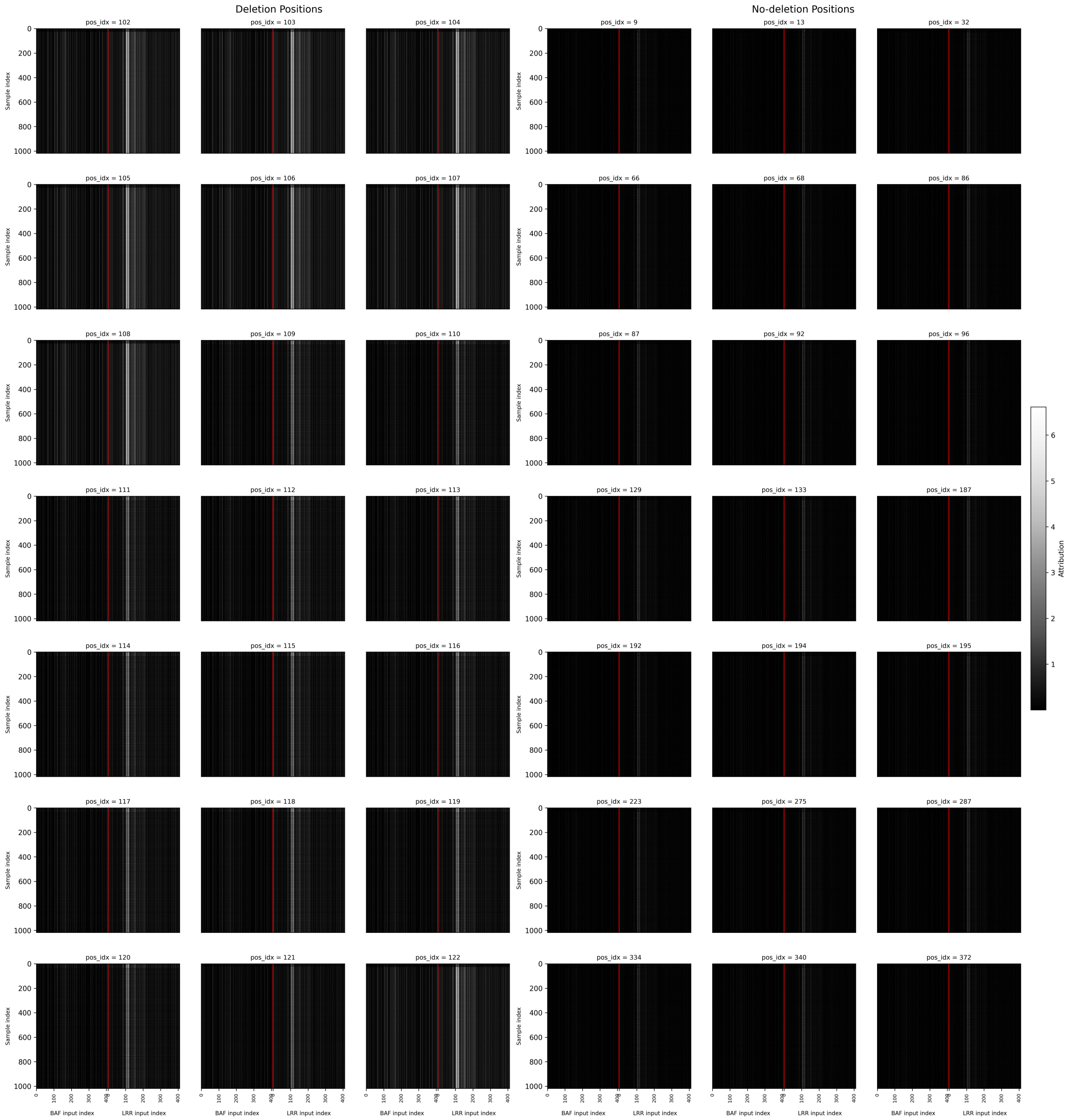


**S5 Fig.** Gradient-based feature attribution heatmaps of nnCNV for each deletion probe (left 3 columns) and randomly selected probes with no deletion (right 3 columns), showing attribution scores across input feature positions (x-axis) for individual samples (y-axis). Vertical red lines separate BAF and LRR input indices.

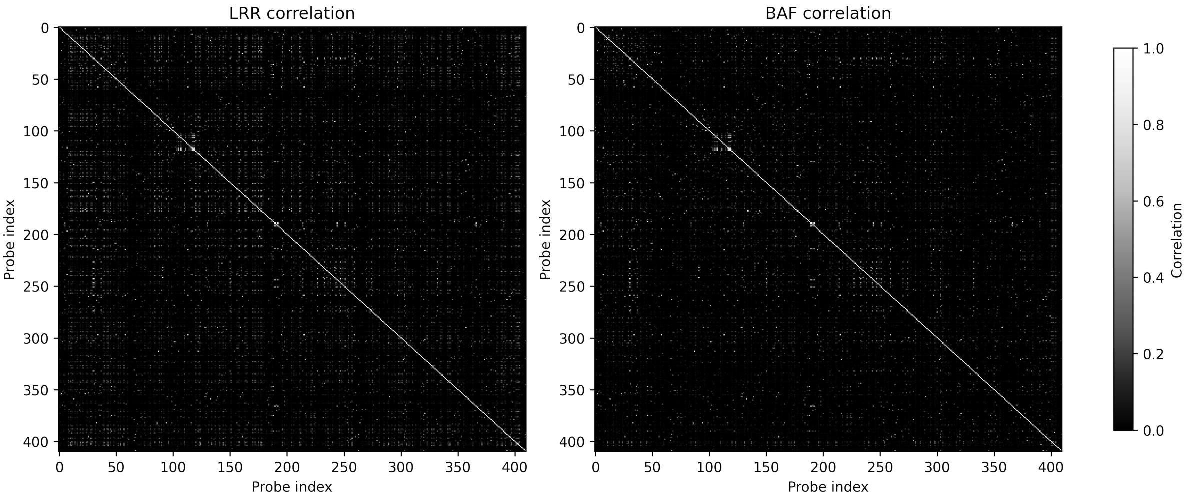


**S6 Fig.** Heatmaps of absolute Pearson's correlation coefficient for LRR (left) and BAF (right) values between positions using combined training, validation and test samples.
